# Rhizoxin and 2,4-diacetylphloroglucinol Contribute to Biocontrol of *Pseudomonas protegens* Pf-5 Against Pea Ascochyta Blight Pathogen *Didymella pinodes*

**DOI:** 10.64898/2026.04.21.719965

**Authors:** Jepri Agung Priyanto, Chiseche Mwanza, Maria Purnamasari, Xiaogang Wu, Li Huang, Qing Yan

**Affiliations:** Department of Plant Sciences and Plant Pathology, Montana State University, Bozeman, Montana, USA; Division of Microbiology, Department of Biology, Faculty of Mathematics and Natural Sciences, IPB University, Bogor, Indonesia; Guangxi Key Laboratory of Agro-Environment and Agro-Product Safety/College of Agriculture, Guangxi University, Nanning 530004, China

**Keywords:** Antibiotics, Biocontrol, DAPG, *Didymella pinodes*, *Paenibacillus*, *Pseudomonas*, Rhizoxin

## Abstract

Biological control using beneficial bacteria is a promising strategy for managing pea Ascochyta blight (AB), yet the underlying mechanisms remain poorly understood. In this study, we identified ten bacterial strains from four genera, including *Bacillus*, *Paenibacillus*, *Peribacillus*, and *Pseudomonas*, that significantly reduced the severity of AB caused by *Didymella pinodes* under greenhouse conditions. Most strains inhibited *D. pinodes in vitro*, suggesting antibiosis as a primary mode of action. To further elucidate the biocontrol mechanisms, we used *Pseudomonas protegens* Pf-5, which produces eight known antimicrobial compounds, as a model. While wild-type Pf-5 strongly inhibited *D. pinodes* in cultures and controlled AB *in planta*, a derivative (Δ8-fold mutant) lacking all eight compounds showed significantly compromised biocontrol efficacy. Individual complementation of biosynthetic genes for rhizoxin, 2,4-diacetylphloroglucinol (DAPG), pyrrolnitrin, or hydrogen cyanide partially restored inhibitory activity, confirming their roles in inhibition of *D. pinodes*. Notably, restoring rhizoxin and DAPG biosynthesis recovered the disease control capability of the Δ8-fold mutant in greenhouse trials. These results demonstrate that rhizoxin and DAPG are key metabolites driving the biocontrol activity of *P. protegens* against *D. pinodes*.

**SIGNIFICANCE:** An advanced understanding of how beneficial bacteria control plant diseases can help us better use these microorganisms in agriculture. In this study, beneficial bacteria isolated from pea roots and soils effectively mitigated damages of pea Ascochyta blight caused by the fungal pathogen *Didymella pinodes*. Most of the identified beneficial bacteria inhibited the fungal pathogen in cultures, indicating antimicrobial compounds were likely produced by the bacteria to control the disease. Using the soil beneficial bacterium *Pseudomonas protegens* Pf-5 as a model, we demonstrated that four bacteria-derived antimicrobial compounds, rhizoxin and 2,4-diacetylphloroglucinol (DAPG), pyrrolnitrin, and hydrogen cyanide play important roles in inhibiting *D. pinodes* growth. This study also showed that rhizoxin and DAPG produced by Pf-5 contribute to the suppression of AB development. These findings provided new insights into the molecular basis of beneficial bacteria-mediated disease suppression of pea Ascochyta blight.

## 1. INTRODUCTION

Biocontrol using beneficial bacteria is promising in plant disease management. However, application of beneficial bacteria is often hampered by varied efficacy *in planta*. Identification of bacteria that have consistent disease control efficacy is critical, and an advanced understanding of their disease control mechanisms can help us better use the beneficial bacteria to control plant diseases.

Pea Ascochyta blight (AB) is a foliar disease caused by several fungal pathogens including *Didymella pinodes* (synonym: *Ascochyta pinodes*, *Peyronellaea pinodes*), *D. pinodella*, *D. pisi*, and *Ascochyta koolunga* (Ahmed et al. 2015; Keirnan et al. 2021; Liu et al. 2023; Davidson et al. 2013). These pathogens can infect pea plants individually or coexist as a complex within a single field or plant (Le May et al. 2009). Among these pathogens, *D. pinodes* has a broad range of hosts including pea (*Pisum sativum*) and at least 19 other legume species (Barilli et al. 2016). *D. pinodes* was reported as the primary pathogen of AB in many pea-growing countries, including Australia (Davidson et al. 2011), Canada (Habibi et al. 2016), China (Liu et al. 2016), France (Le May et al. 2018) and United States (Fosenka et al. 2023; Owati et al. 2020),

Several bacterial strains have been identified to control pea AB. However, the mechanisms of how beneficial bacteria control pea AB remain largely unknown. For example, *Pantoea agglomerans*, *Bacillus amyloliquifaciens* and *Bacillus subtilis* significantly reduced pea AB disease severity (Liu et al. 2016). These bacteria inhibited *D. pinodes* in cultures, although the antimicrobial compound(s) were not identified and their roles in disease control were not determined (Liu et al. 2016). Additionally, application of *Bacillus megaterium* and *Pseudomonas fluorescens* reduced pea AB disease severity, but the mechanisms remain unclear (Ahmed & Agha 2022).

Pea AB has been reported in Montana (Owati et al. 2020), highlighting the need for effective disease management strategies. Indigenous bacteria are often well adapted to local abiotic and biotic conditions (Kumar & Gopal 2015), enabling them to compete with pathogens and other microorganisms for nutrients and spaces. Previous studies have demonstrated the effectiveness of indigenous bacteria in controlling plant diseases (Pérez-Rodríguez et al. 2020; Reddy et al. 2024; Castillo-Novales et al. 2025). Isolating and characterizing beneficial bacteria from pea farms in Montana is expected to yield effective biocontrol agents to control pea AB.

The first goal of this study was to identify beneficial bacteria that control pea AB caused by *D. pinodes*. To achieve this goal, bacterial strains were isolated from pea tissues and soils collected in Montana. Beneficial bacteria that strongly reduced AB disease severity were identified. The second goal was to investigate the mechanisms used by the beneficial bacteria to control pea AB. To this end, *Pseudomonas protegens* Pf-5 was used because it has been served as a model bacterium in studies of biocontrol mechanisms (Brodhagen et al. 2004; Kidarsa et al. 2013; Lai et al. 2022). Pf-5 produces at least eight antimicrobial compounds, including pyrrolnitrin (Howell & Stipanovic 1979), hydrogen cyanide (Kraus & Loper 1992), 2,4-diacetylphloroglucinol (DAPG) (Nowak-Thompson et al. 1994), pyoluteorin (Howell & Stipanovic 1980; Nowak-Thompson et al. 1999), orfamide A (Gross et al. 2007), rhizoxin (Brendel et al. 2007; Loper et al. 2008), toxoflavin (Philmus et al. 2015), and polyyne (Hotter et al. 2021). Production of these compounds is activated by a global regulator GacA in Pf-5 (Kidarsa et al. 2013, Philmus et al. 2015, Purnamasari et al. 2025, Mwanza et al. 2025). Previous research indicated that Pf-5 could control pea AB although the underlying mechanism was not investigated (Annan et al. 2023). In this study, the antimicrobial compounds required for Pf-5 to inhibit *D. pinodes* were determined and their roles in pea AB control were characterized.

## 2. MATERIALS AND METHODS

### 2.1 Microbial Materials and Growth Conditions

*D. pinodes* isolate C6 was isolated previously from Ascochyta blight (AB)-contaminated pea seeds (Annan et al. 2023). To promote conidia production, the isolate was routinely cultured on pea meal agar (PMA) at 28°C, prepared by combining 25 g of blended dry pea seeds and 15 g of agar (BD Difco, Becton Dickinson and Company) in 1 L of MiliQ water.

Biocontrol bacteria used in this study include the ones isolated in this work and the model strain *P. protegens* Pf-5 (Howell & Stipanovic 1979). To isolate pea-associated bacteria, pea leaves, stems, roots and rhizosphere soils were collected from pea farms in Montana. One gram of the collected samples was homogenized in 10 mL sterilize MiliQ water. A 100-µL aliquot of the suspension was spread onto various culture media, including Reasoner’s 2A agar (R2A, Neogen), potato dextrose agar (PDA, BD Difco™, Becton Dickinson and Company), and tryptic soy agar (TSA, BD Difco™, Becton Dickinson and Company). After 48 h of incubation at 28°C, colonies with different morphologies were purified and stored at −80°C for further study.

*P. protegens* Pf-5 was originally isolated from cotton field soils (Howell & Stipanovic 1979). Pf-5 and its derivatives used in this study were listed in the Table 1. The bacterium was cultured at 28°C in King’s B medium (KB) (King et al. 1954) unless other specific media were described.

**Table 1:**
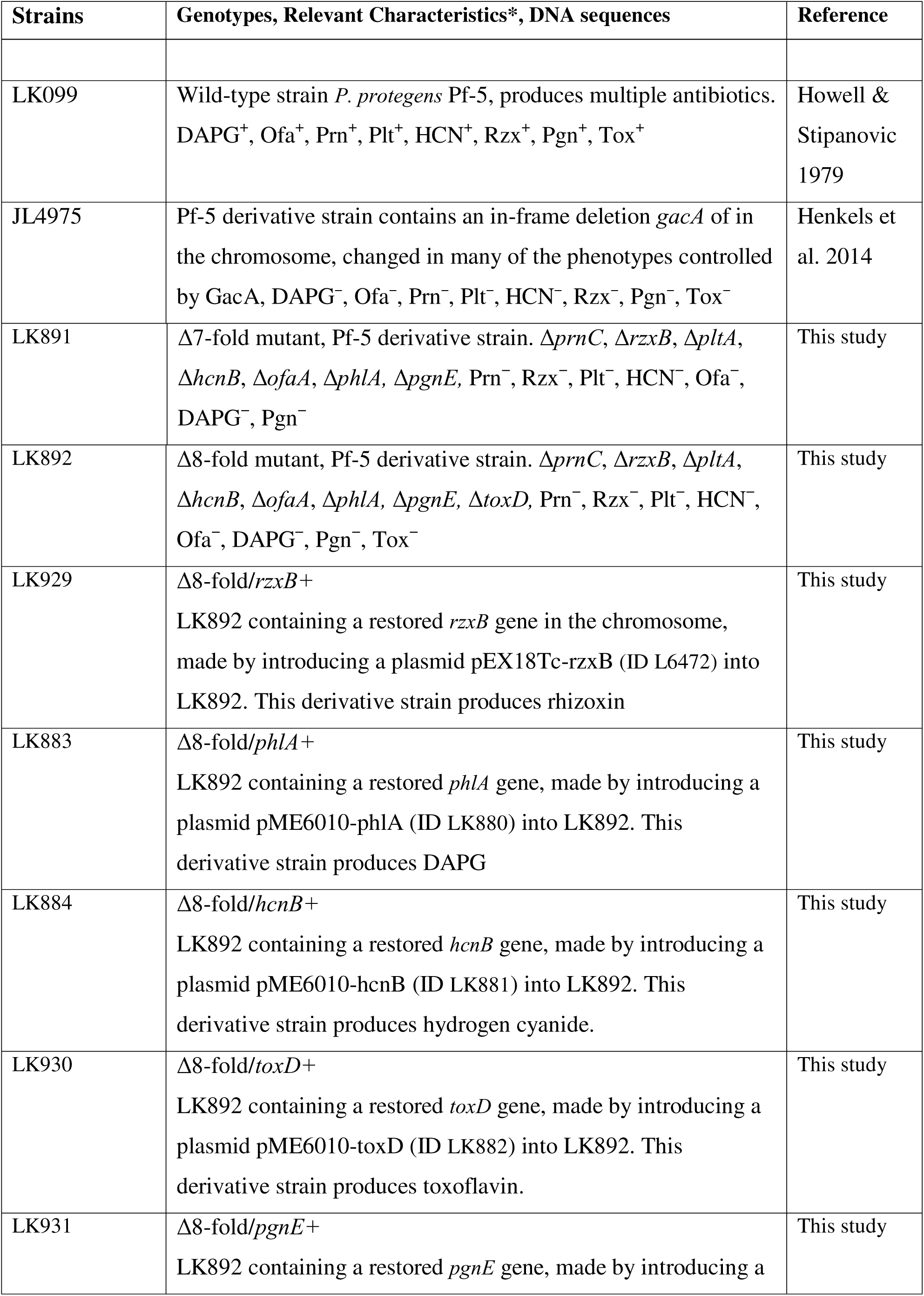
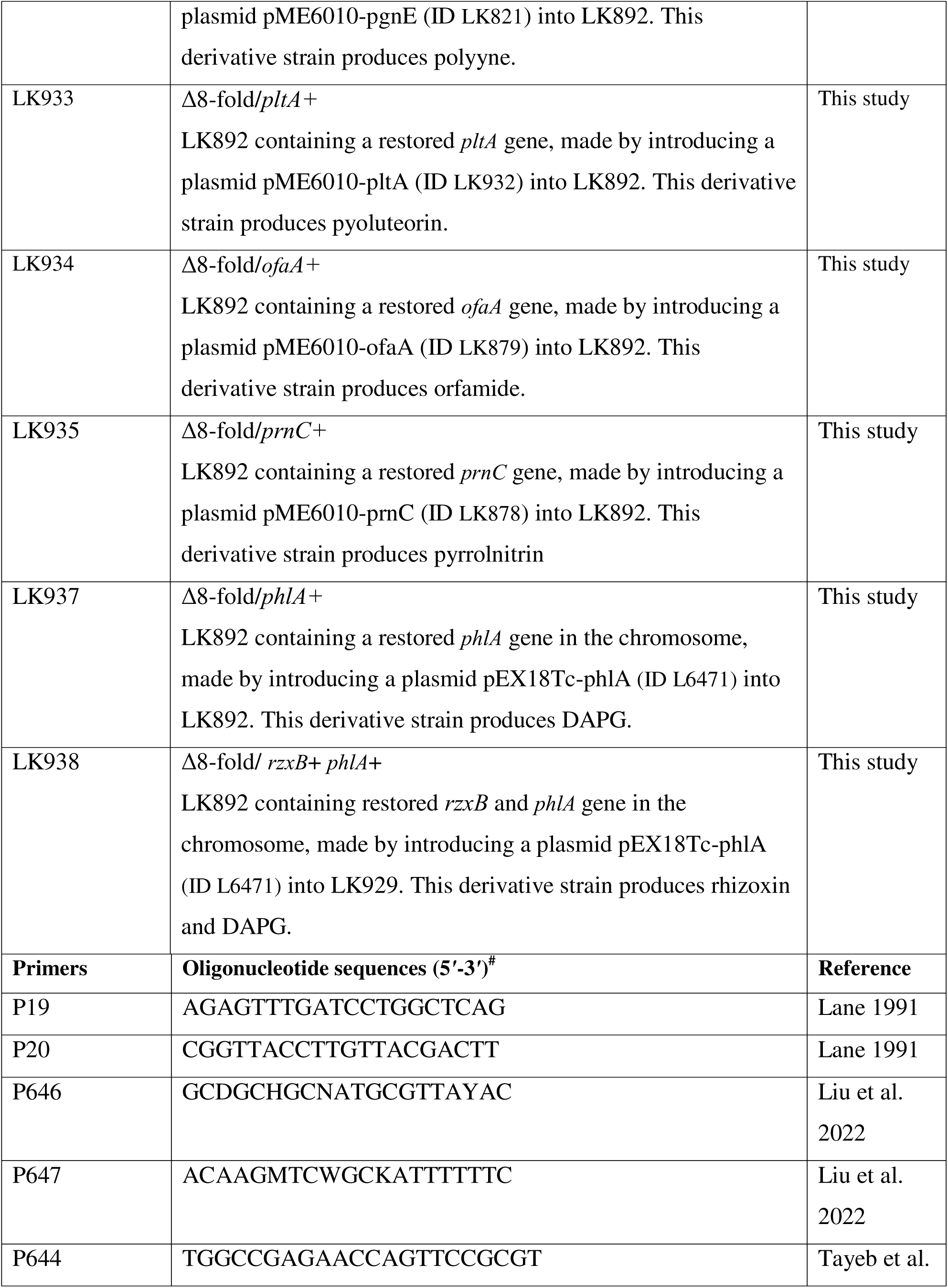
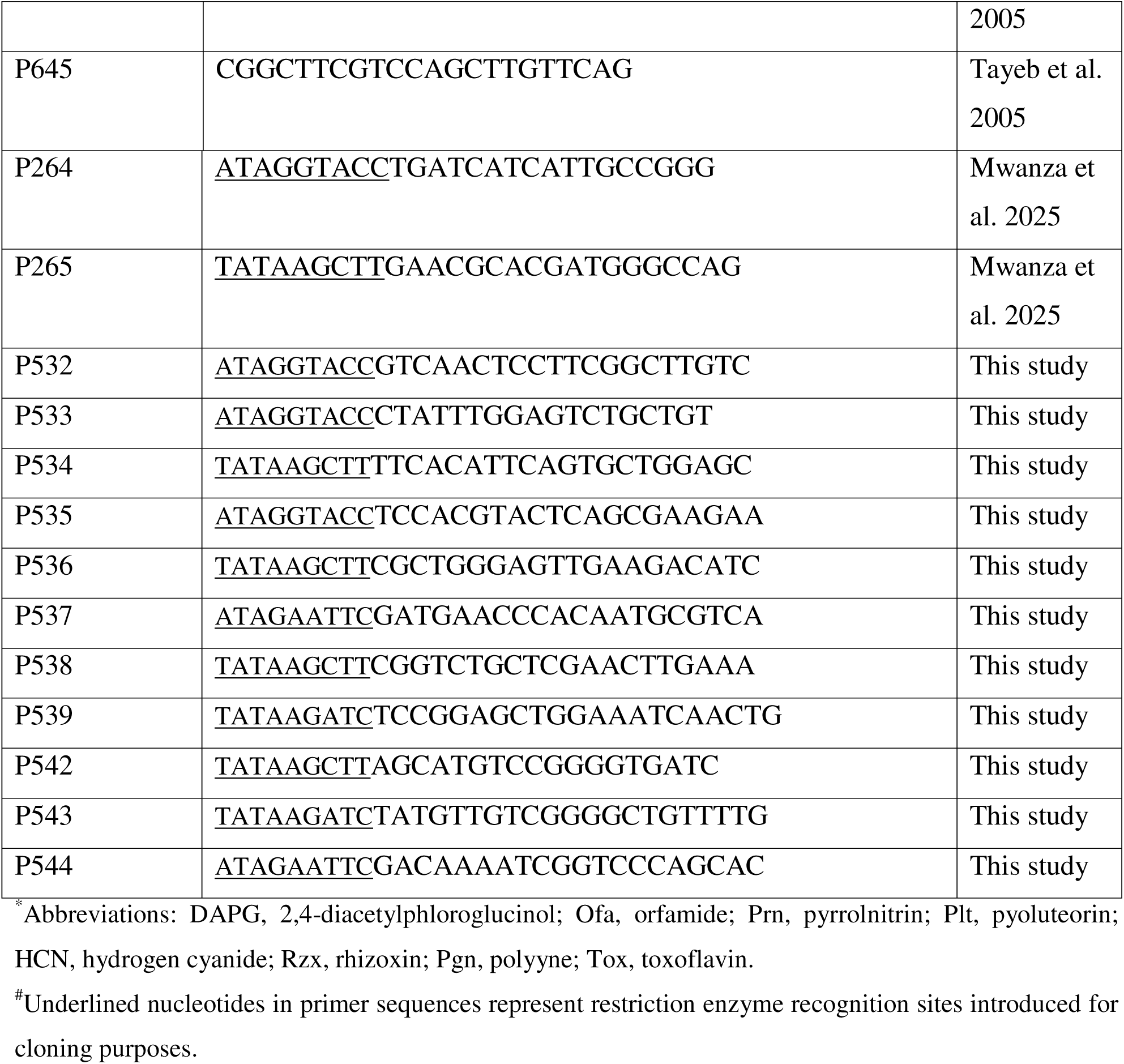
Bacterial Strains and Primers.

### 2.2 Plant Materials and Growth Conditions

The seeds of pea (*Pisum sativum*) variety Fallon were obtained from the Montana State University (MSU) Pulse Breeding Program. The seeds were surface-sterilized with 3% sodium hypochlorite for 2 minutes, rinsed three times with sterile water, and air-dried prior to sowing in Sunshine soil (Sun Gro Horticulture, Agawam, MA). The plants were then grown in the greenhouse at the MSU Plant Growth Centre under controlled conditions, including a day/night temperature of 30.2/18°C, 22% relative humidity, and a 16-h photoperiod. After two weeks of growth, the plants were considered ready for inoculation.

### 2.3 Biocontrol Assays

#### 2.3.1 Preparation and Inoculation of Beneficial Bacteria and Fungal Pathogen

Bacteria strains were cultured overnight at 28°C on Luria–Bertani (LB, BD Difco™, Becton Dickinson and Company) plates or KB plates. Cells were harvested from the plates and washed three times with sterile water. Approximately 20 mL of each bacterial suspension (OD_600_ = 1.0), amended with one drop of Tween 20, was spray-inoculated onto leaves of the two-week-old pea plants until the leaf surfaces were uniformly coated and visibly wet. The plants were maintained in a dew chamber at 22°C for 24 hours prior to *D. pinodes* inoculation.

A mixed inoculum of the beneficial bacteria was prepared by mixing *P. protegens* Pf-5 and the individual pea-associated bacterial isolate, yielding 20 mL of a mixed bacterial suspension (final cell density at OD□□□ = 1.0 for both strains). The mixed inoculum was spray-inoculated onto pea leaves as described above.

To prepare the pathogen inoculum, *D. pinodes* was cultured on PMA plates for two weeks at 28°C. The culture plate was flooded with 30 mL of sterile distilled water and gently scraped with a glass rod to dislodge conidia. The spore suspension was filtered through a double layer of sterilized cheesecloth, adjusted to 1 × 10^5^ spores/mL in 200 mL of water supplemented with two drops of Tween 20, then spray-inoculated onto the pea leaves. The pea plants were placed in a dew chamber at 22°C in darkness for 24 hours before being transferred to a greenhouse and incubated in a plastic tent to maintain a high humidity. Plants sprayed only with sterile water or the pathogen served as negative and positive controls, respectively. Each treatment consisted of 10 pots with one seed per pot. The experiment was repeated independently three times.

#### 2.3.2 Disease Evaluation

The AB disease was evaluated at seven days after pathogen inoculation. The severity of symptoms on leaves, stems, and stipules was assessed using a Disease Severity Index (DSI) including a scale of the disease severity between 0 to 5 as shown in Figure 1A and described below:

0 = No visible lesions on any part of the plant; plant appears healthy.
1 = A few small leaf flecks.
2 = Less than 25% of the plant leaves show lesions; the stem is unaffected.
3 = Numerous flecks or lesions on leaves and/or stems; less than 50% coverage.
4 = Large, coalescing lesions covering more than 50% of the plant surfaces; plant still alive.
5 = Severe lesions with stem girdling; plant is dead.

**Figure 1.**
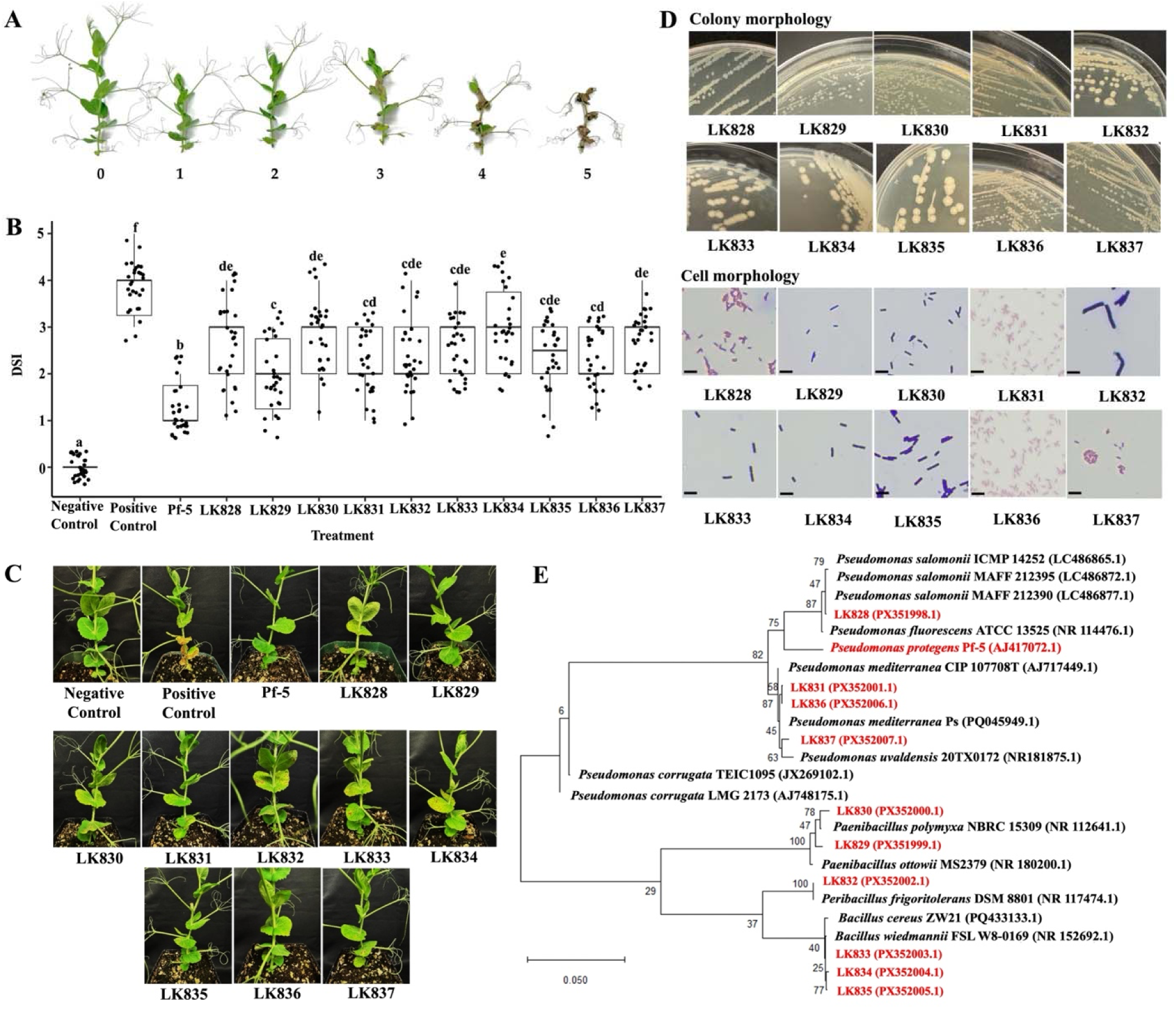
Biocontrol activity of *P. protegens* Pf-5 and pea-associated bacteria against pea Ascochyta blight. **A.** Pea plants representing each level of the standard Disease Severity Index (DSI). **B.** Disease severity of pea plants treated with *P. protegens* Pf-5 and the pea-associated bacteria isolates ten days after inoculation of *D. pinodes*. Data were presented as mean ± standard deviation. Each dot represents an individual plant. A total of 30 plants were used in one treatment. The experiment was repeated independently three times. Different letters indicate significant differences among treatments (ANOVA, Tukey’s test, *p* < 0.05). **C.** Results of the biocontrol assay. Photos of the representative plant were taken at seven days after fungal inoculation. **D.** Colony and cell morphology of the bacterial isolates grown on LB medium at 28 °C for 24 h. A cell with a pink or purple colour indicates the isolate was identified to be Gram-negative or Gram-positive, respectively, via the Gram staining assay. The bar size is 10 µm. **E.** Phylogenetic tree of the tested bacterial strains (highlighted in red) constructed via multiple genes (16S rRNA, *rpoB*, *gyrA*) sequences analysis. Numbers at the node indicate the percentage of bootstrap support based on a maximum likelihood method of 1,000 resampled data sets. The scale bar represents 0.05 substitution per nucleotide position.

#### 2.3.3 Disease Control Assay to Test Duration of the Beneficial Bacteria

Pea leaves were spray-inoculated with bacterial suspensions (OD□□□= 1) until evenly wetted, then incubated for 24 hours in a dew chamber. Subsequently, *D. pinodes* spores were applied at three time points post-bacterial application: 1, 7, and 14 days. DSI was measured seven days after pathogen inoculation. The bacterial survival on leaves was also measured: inoculated leaves were homogenized in sterile water, and the bacterial populations were quantified by spreading a serial dilutions of the bacterial suspension on KB plates supplemented with 100 µg/mL streptomycin for Pf-5 and on LB plates supplemented with 100 µg/mL ampicillin for LK829. Bacterial populations were expressed as CFU/g of leaf tissues.

### 2.4 Inhibition Assays of Selected Isolates against *D. pinodes*

The inhibition of beneficial bacteria against *D. pinodes* was evaluated using two different methods: mycelium-based inhibition assay and spore-based inhibition assay, as described below.

#### 2.4.1 Mycelium-Based Inhibition Assay

A 6-mm mycelial plug from a seven-day-old *D. pinodes* culture was placed at the center of a PMA plate and incubated at 28°C for 7 days. Subsequently, 3 µL of bacterial suspension OD□□□ = 1 was spotted 2.5 cm from the edge of the fungal colony. The plates were allowed to air-dry and then incubated for an additional 7 days at 28°C before the inhibition zone was measured.

#### 2.4.2 Spore-Based Inhibition Assay

Spores from a two-week-old *D. pinodes* culture were harvested by flooding the culture with sterile water and filtering the suspension through six layers of sterile Miracloth (Millipore Sigma). A 200-µL aliquot of the spore suspension (1 × 10^5^ spores/mL) was added to 20 mL of molten PDA (50°C), resulting in a final concentration of 1 × 10^3^ spores/mL. The mixture was then gently poured into Petri dishes. Once the medium solidified, 3 µL of bacterial suspension (OD_600_ = 3) was spotted onto the agar surface and incubated at 28°C for 3 days. Using the same procedure, inhibitory activity was also evaluated on pea leaf agar (PLA). PLA was prepared by homogenizing 10 g of fresh pea leaves in 100 mL of distilled water, followed by filtration through two layers of Miracloth (MilliporeSigma) to obtain a leaf extract. A total of 100 mL of this filtrate was then mixed with 100 mL of distilled water containing 1 g of blended dry pea seeds. The medium was subsequently autoclaved prior to use.

The results of both assays were evaluated by measuring the inhibition zone between the edge of the bacterial colony and the fungal colony. The assays were performed in three independent experiments, each with two replicates.

### 2.5 Taxonomic Characterization of Beneficial Bacterial Isolates

Bacterial genomic DNA was extracted using Monarch Spin gDNA Extraction kit (New England Biolabs). All bacterial isolates were initially identified based on their 16S rRNA gene sequences by following a previous study (Lane 1991). Isolates of *Pseudomonas* spp. and *Bacillus* spp. were further analysed by sequencing *rpoB* and *gyrA*, respectively. Each gene was amplified by PCR using primer pairs 16Sr RNA (P19/P20), *rpoB* (P644/p645), *gyrA* (P646/P647) as developed by previous studies (Lane 1991; Liu et al 2022; Tayeb et al. 2005) (Table 1). PCR amplification was carried out in a 50-µL reaction mixture containing 2 µL of genomic DNA (100 ng/µL), 2 µL of each primer, 25 µL of KOD 2× Master Mix, and 19 µL of deionized water. PCR conditions were as follows: initial denaturation at 95°C for 5 min; followed by 35 cycles of denaturation at 95°C for 30 s, annealing at 55°C (16S rRNA), 50°C (*gyrA*) or 54°C (*rpoB*) for 30 s, and extension at 72°C for 40 s; with a final extension at 72°C for 5 min. The expected amplicon sizes for 16S rRNA, *gyrA,* and *rpoB* are 1492, 500 and 1247 bp, respectively. All the sequences were deposited to NCBI GenBank and can be accessed via the accession numbers listed in Figure 1E. Multigene alignment was performed using the MUSCLE algorithm, and the phylogenetic tree was constructed using the maximum likelihood method in Molecular Evolutionary Genetics Analysis (MEGA) version 12.1.

### 2.6 Construction of *P. protegens* Pf-5 Derivatives

#### 2.6.1 Construction of Pf-5 Mutants

The Δ7-fold mutant (Δ*pltA*Δ*prnC*Δ*ofaA*Δ*rzxB*Δ*hcnB*Δ*phlA*Δ*toxD*, strain LK891) was made by deleting *toxD,* using a deletion construct pEX18Tc-Δ*toxD* (Philmus et al. 2015), from the chromosome of a Δ6-fold mutant (Δ*pltA*Δ*prnC*Δ*ofaA*Δ*rzxB*Δ*hcnB*Δ*phlA*, strain LK147) (Loper et al. 2016). Next, the Δ8-fold mutant (Δ*pltA*Δ*prnC*Δ*ofaA*Δ*rzxB*Δ*hcnB*Δ*phlA*Δ*toxD*Δ*pgnE*, strain LK892) was made by deleting *pgnE,* using a deletion construct pEX18Tc-Δ*pgnE* (Mwanza et al. 2025), from the chromosome of LK891. The mutants were generated following our previous protocol (Yan et al. 2021). Briefly, the deletion constructs were introduced into Pf-5 cells via electroporation (1.8 kV, 5 ms). Single-crossover recombinants were selected on KB amended with tetracycline, followed by selection of double-crossover recombinants on LB amended with sucrose to resolve the plasmid backbone using the SacB-based counterselection. The deletions of target genes were confirmed by whole sequencing analysis as described below.

#### 2.6.2 Whole Genome Sequencing of Pf-5 Mutant to Validate Mutations

The genome of the Δ8-fold mutant (strain LK892) was sequenced to confirm the mutation of target genes for the eight antimicrobial compounds. Genomic DNA was extracted using the MasterPure Complete DNA and RNA Purification Kit (Lucigen, Middleton, WI). Long-read whole-genome sequencing was performed on an Oxford Nanopore MinION platform using a standard ligation-based library preparation workflow and ONT flow cell chemistry. Raw reads were basecalled with ONT basecalling software, and adapter/low-quality sequences were removed prior to assembly. A total of 532,268 raw reads were generated and used for genome assembly on the Tempest high-performance computing cluster using a standard long-read assembly pipeline, followed by polishing to improve consensus accuracy. The final assembly was annotated with Bakta, and circular genome maps (CDS, RNA features, GC content, and GC skew) were generated using Proksee (Grant et al. 2023).

To determine the mutation of the target eight genes, sequences spanning each candidate antibiotic biosynthetic gene were extracted from the assembled mutant genome and compared to the genome of the wild-type Pf-5 using NCBI BLAST searches. BLAST alignments were used to identify the corresponding Pf-5 orthologs and to define the mutations in the Δ8-fold mutant (strain LK892). Coordinates from the Pf-5 reference were used to report mutation boundaries and to generate locus schematics for the eight antibiotic-associated genes. To confirm that the Δ8-fold mutant genotype was otherwise conserved, we performed a genome-wide comparison against the Pf-5 wild-type reference and screened the variant list for additional high-confidence disruptive changes; no other mutations with clear predicted loss-of-function effects were detected beyond the eight antibiotic-associated loci.

#### 2.6.2 Construction of Complemented Strains of Pf-5

Individual complementation of seven antibiotic biosynthetic genes (*phlA*, *prnC*, *toxD*, *hcnB*, *ofaA*, *pltA*, *pgnE*) was made by introducing the corresponding wild-type genes, carried by pME6010 vector, into the Δ8-fold mutant. Each of the genes was amplified by PCR using the following primer pairs: *phlA* (P533/P534), *prnC* (P531/P532), *toxD* (P535/P536), *hcnB* (P537/P538), *ofaA* (P539/P542), *pltA* (P543/P544), and *pgnE* (P264/P265). The constructs were verified by sequencing and introduced into LK892 by electroporation (1.8 kv, 5 ms). The double complementation of *phlA* and *rzxB* in the Δ8-fold mutant was made through chromosomal replacement using pEX18Tc-based constructs containing the wild-type *phlA* (Kidarsa et al. 2011) or *rzxB* (Henkels et al. 2014). The chromosomal replacement was performed following the above method of gene deletion. The complementation was confirmed by PCR analysis and phenotype assays as described below.

Restored production of the eight antimicrobial compounds in the Δ8-fold mutant was further validated by phenotypic assays, including direct visualization of the compounds via thin-layer chromatography (TLC), inhibition against microorganisms sensitive to the compounds, and other bacterial traits that rely on the compounds. Specifically, Δ8-fold/*rzxB*^+^ was tested via inhibition of *D. pinodes* which is sensitive to rhizoxin (this study); Δ8-fold/*phlA*^+^ was tested via inhibition of *Aphanomyces euteiches* which is sensitive to DAPG (Lai et al. 2022); Δ8-fold/*toxD*^+^ and Δ8-fold/*pgnE*^+^ were tested via inhibition of *Pseudomonas syringae* which is sensitive to toxoflavin and polyyne (Philmus et al. 2015; Mwanza et al. 2025); Δ8-fold/*ofaA*^+^ was tested via a swarming motility assay on 0.6% agar M9 minimal medium because orfamide is needed in bacterial motility (Lim et al. 2012); Δ8-fold/*hcnB*^+^ was tested via a picric acid-based colorimetric assay (Torrescassana et al. 2025); Strains Δ8-fold/*pltA*^+^ (pyoluteorin-producing) and Δ8-fold/*prnC*^+^ (pyrrolnitrin-producing) were cultured in nutrient broth supplemented with 1% glycerol for 24 h and on pea meal agar plates for 7 days, respectively. Extracts obtained with ethyl acetate were subsequently analyzed by TLC to confirm metabolite production (Quecine et al. 2016).

### 2.7 Statistical Analysis

Statistical analysis was performed using R software (version 4.5.3). One-way ANOVA was used for single-factor experiments, followed by Tukey’s HSD test for post hoc comparisons at *p* < 0.05.

## 3. RESULTS

### 3.1 Identification of Bacterial Isolates with Biocontrol Activity

A total of 350 bacterial strains were isolated from pea rhizosphere soils, leaves, and stems and screened for their ability to control pea AB caused by *D. pinodes* in the greenhouse. The ten most effective isolates (LK828 - LK837), that showed significant and consistent biocontrol activity in at least three independent experiments, were selected for further investigation (Fig. 1B, 1C). Gram staining and microscopic analyses revealed that four isolates were Gram-negative and six were Gram-positive (Fig. 1D). Phylogenetic analysis based on sequence alignment of three housekeeping genes (16S rRNA, *rpoB*, and *gyrA*) showed that four isolates (LK828, LK831, LK836, LK837) belonged to *Pseudomonas*, three (LK833, LK834, LK835) to *Bacillus*, two (LK829, LK830) to *Paenibacillus*, one (LK832) to *Peribacillus* (Fig. 1E).

This study also includes a soil bacterium *P. protegens* Pf-5 because of its previously reported biocontrol activity against pea AB (Annan et al. 2025). Our result shows that Pf-5 exhibited a stronger biocontrol activity than the ten pea bacterial isolates (Fig. 1B, 1C).

### 3.2 Beneficial Bacteria Protected Pea Plant from *D. pinodes* for Up to Two Weeks

*P. protegens* Pf-5 and *Bacillus* sp. LK829, the most effective Gram-negative and Gram-positive bacterial isolates identified in this study, were further characterized for their biocontrol traits. To evaluate the duration of protection against *D. pinodes*, pea plants treated with the beneficial bacteria were kept in a greenhouse for 1, 7, or 14 days prior to pathogen inoculation. A single application of Pf-5 protected pea plants for at least 14 days (Fig. 2A). Pf-5 exhibited the highest biocontrol efficacy immediately after application on pea leaves, with efficacy gradually declining over time. To assess the persistence of Pf-5 on plant, its population density on pea leaves was quantified at multiple time points. The population declined from an initial density of 2.5 × 10^8^ CFU/g to 1.3 × 10^5^ CFU/g by day 14 (Fig. 2B).

**Figure 2.**
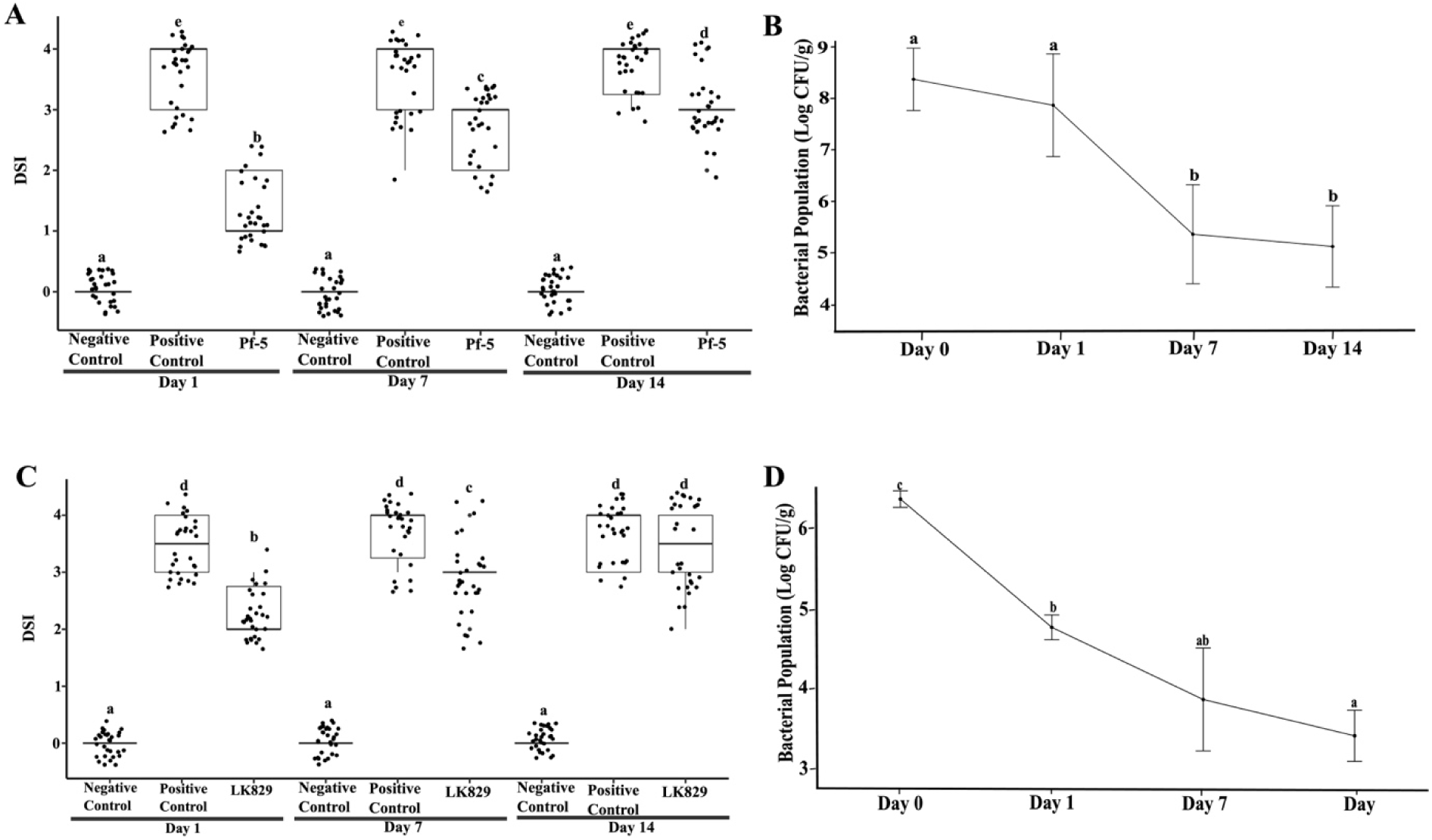
Biocontrol activity against *D. pinodes* at 1, 7, and 14 days after bacterial application. **A.** Disease severity index (DSI) of pea plants inoculated with *P. protegens* Pf-5. Each treatment comprised 10 replicates, and the experiment was independently repeated three times. Data were presented as means ± standard deviation. Different letters indicate significant differences among treatments (ANOVA, Tukey’s test, *p* < 0.05). **B.** Population dynamics of Pf-5 on pea leaves. **C.** DSI of pea plants treated with LK829. **D.** LK829 population dynamics on pea leaves. Leaves inoculated with the bacteria were sampled at 0, 1, 7, and 14 days after bacterial application. Bacterial populations were quantified using the serial dilution method and expressed as Log CFU/g.

A similar disease control profile was observed for strain LK829, which protected pea plants from *D. pinodes* infection for up to 7 days following plant application (Fig. 2C). The population density of LK829 on pea leaves gradually declined from an initial level of 2.5 × 10^6^ CFU/g to 2.5 × 10^3^ CFU/g by day 14 (Fig. 2D).

### 3.3 Mixed Inoculation of Different Beneficial Bacteria Did Not Enhance Biocontrol Efficacy

To determine whether different beneficial strains could interact synergistically to enhance disease control, Pf□5 was individually mixed with each of the selected pea associated beneficial bacteria and sprayed onto pea leaves prior to inoculation with *D. pinodes*. Overall, co□inoculation of Pf□5 with any of the ten pea associated strains did not improve biocontrol efficacy compared with Pf□5 applied alone (Fig. 3). In fact, co□inoculation with four strains (LK828, LK829, LK836, and LK837) reduced the biocontrol activity of Pf□5, whereas mixtures with the remaining six strains resulted in disease control levels comparable to Pf□5 single inoculation.

**Figure 3.**
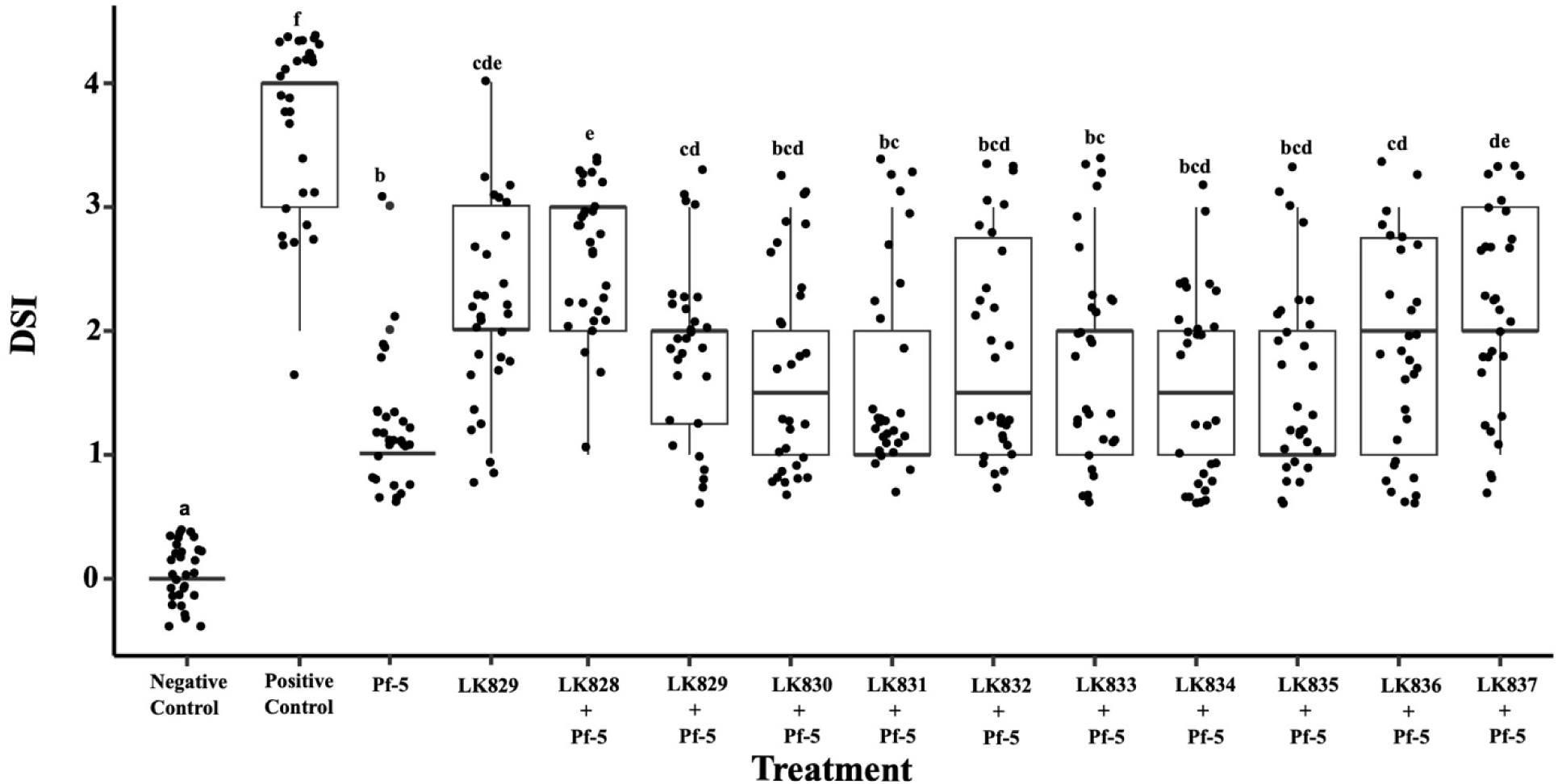
Biocontrol activity of mixed inoculation of Pf-5 with the pea-associated bacteria. A single application of Pf-5 and LK829 were also included in the assay. The disease severity index (DSI) of each treatment was shown. A total of 30 plants were used in each treatment. The experiment was repeated independently three times. Data were presented as means ± standard deviation. Different letters indicate significant differences among treatments (ANOVA, Tukey’s test, *p* < 0.05).

A single inoculation of LK829 was included as a reference to facilitate comparison with its mixed inoculation with Pf□5. The combined application of LK829 and Pf□5 provided greater disease suppression than LK829 alone but was less effective than Pf□5 applied individually.

Despite these differences, all mixed inoculations resulted in significantly reduced disease symptoms compared with the pathogen only control.

### 3.4 Most of the Beneficial Bacterial Isolates Inhibited *D. pinodes* in Cultures

To investigate the mechanism used by the beneficial bacteria to control *D. pinodes*, their antagonistic activity against *D. pinodes* was evaluated using both mycelium-based inhibition assay and spore-based inhibition assay. In the mycelium-based inhibition assay, a plug of *D. pinodes* was inoculated at the centre of the culture plate, while the bacterial strains were inoculated at the periphery. Using this approach, six isolates (Pf-5, LK829, LK830, LK833, LK834, LK835) exhibited inhibitory activity against *D. pinodes* on PMA plates (Fig. 4A).

**Figure 4.**
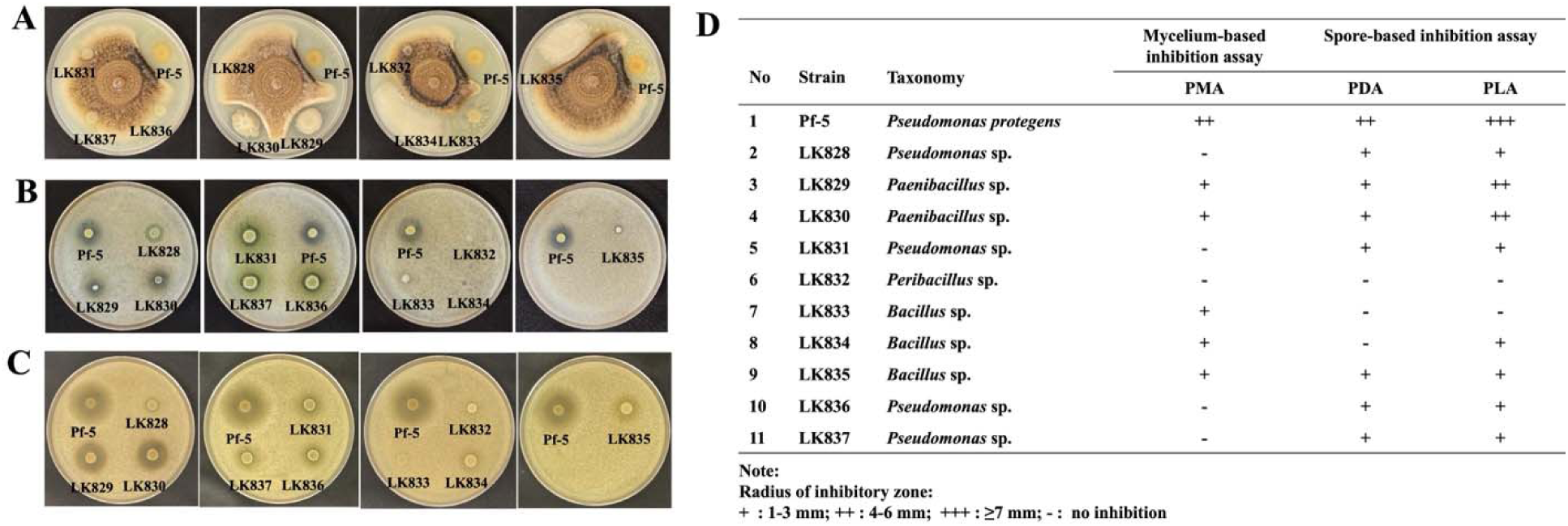
Inhibitory activity of *P. protegens* Pf-5 and pea-associated bacteria against *D. pinodes*, evaluated using different inhibition assays. **A.** Mycelial growth inhibition on pea meal agar (PMA) one week after bacterial inoculation. **B.** Fungal growth inhibition assessed using a spore-based inhibition assay on PDA. **C.** Fungal growth inhibition on pea leaf agar (PLA). **D.** Radius of the inhibitory zone for each strain, expressed in millimetres (mm). Images are representative of two technical replicates, and all experiments were independently repeated three times. Inhibitory activity was scored semi-quantitatively as –, +, ++, and +++, representing no, weak, moderate, and strong inhibition based on the radius of the inhibition zone between the edges of the bacterial and fungal colonies.

Antagonistic activity was further evaluated using a spore-based inhibition assay, in which fungal spores were premixed into molten PDA prior to bacterial inoculation. Using this method, eight isolates inhibited *D. pinodes* on PDA plates (Fig. 4B). To better mimic the nutrient environment of pea leaves, pea leaf agar (PLA) was developed and used in the spore-based inhibition assay. Nine isolates exhibited inhibitory activity on PLA plates (Fig. 4C).

Overall, ten of the eleven tested bacterial strains, except *Paenibacillus* sp. LK832, inhibited *D. pinodes* under at least one culture condition (Fig. 4E). Notably, four isolates (Pf-5, LK835, LK829 and LK830) displayed inhibitory activity in all assays. One isolate, LK834, showed inhibitory activity only on pea-based media (PMA and PLA).

### 3.5 Rhizoxin, DAPG, Pyrrolnitrin and Hydrogen Cyanide Contribute to the Inhibition of *D. pinodes* by *P. protegens* Pf-5

To identify bacterial metabolites responsible for inhibition of *D. pinodes*, *P. protegens* Pf□5 was used as a model because of its strong antagonistic activity (Fig. 4) and well characterized antimicrobial biosynthetic pathways. Deletion of *gacA* abolished Pf□5-mediated inhibition of *D. pinodes* (Fig. 5A), indicating that inhibition depends on GacA regulated antimicrobial compounds. Pf□5 is known to produce at least eight GacA regulated antimicrobial metabolites (Kidarsa et al. 2013, Philmus et al. 2015, Purnamasari et al. 2025, Mwanza et al. 2025). To determine whether these compounds contribute to fungal inhibition, an eight fold deletion mutant lacking all corresponding biosynthetic genes (Δ8□fold mutant; strain LK892) was constructed and confirmed by whole genome sequencing (Fig. 5B, 5C). The Δ8□fold mutant failed to inhibit *D. pinodes* (Fig. 5A), suggesting that at least one of the antimicrobial compounds is required for inhibition.

**Figure 5.**
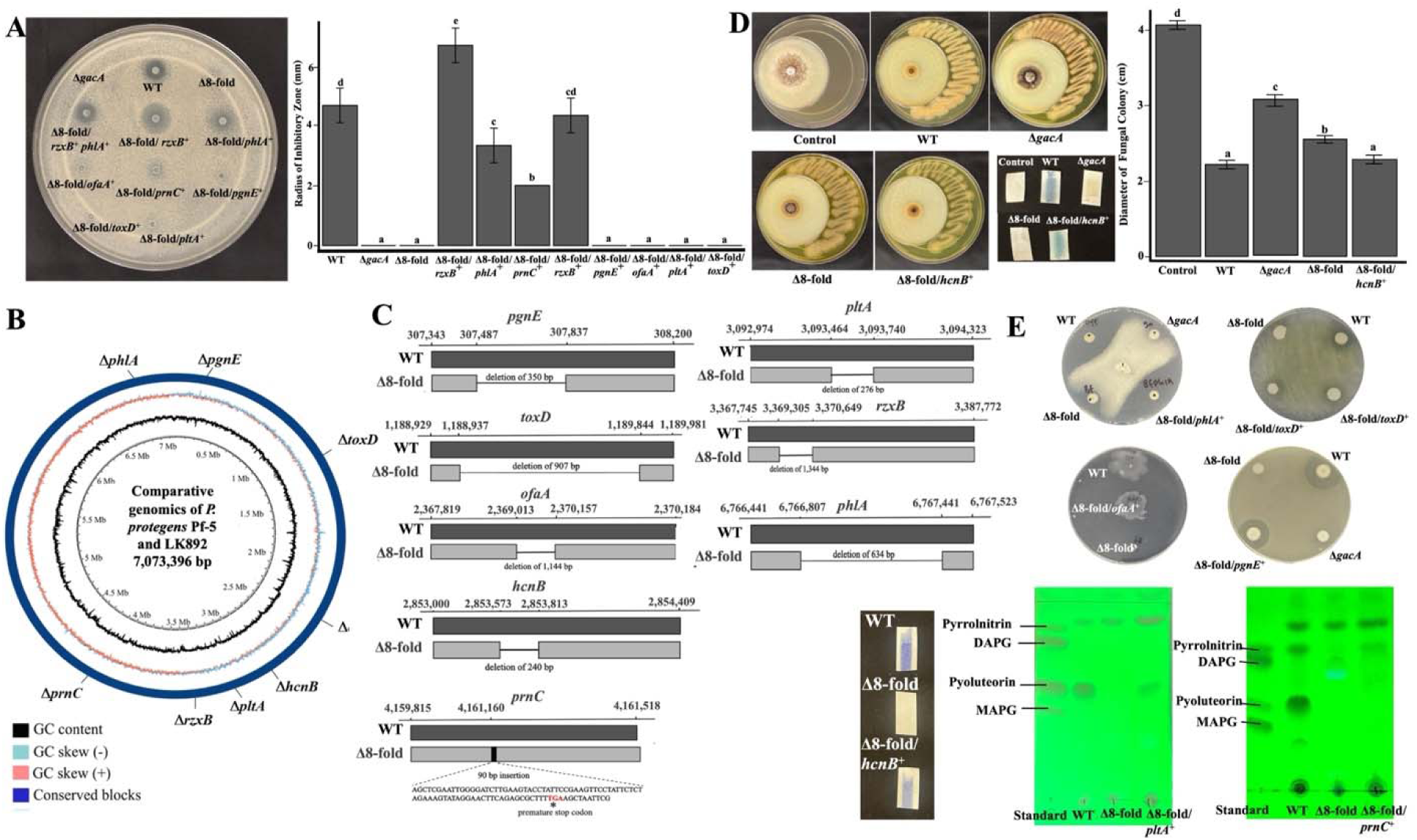
Identification of antimicrobial compounds required for *P. protegens* Pf-5 to inhibit *D. pinodes* in cultures. **A.** Inhibition of Pf-5 and its derivative strains against *D. pinodes* on a PDA plate. Each bacterial suspension (OD□□□ = 1) was spot-inoculated onto PDA medium containing 5 × 10□ spores/mL of *D. pinodes*. Plates were incubated at 28°C for three days before recording the results. WT: wild type; Δ8-fold: Δ*prnC*Δ*rzxB*Δ*pltA*Δ*hcnB*Δ*ofaA*Δ*phlA*Δ*pgnE*Δ*toxC*; Δ8-fold/*rzxB^+^*: 8-fold mutant complemented with *rzxB;* Δ8-fold/*rzxB^+^phlA^+^*: 8-fold mutant complemented with *rzxB* and *phlA;* Δ8-fold/*phlA^+^*: 8-fold mutant complemented with *phlA;* Δ8-fold/*prnC^+^*: 8-fold mutant complemented with *prnC;* Δ8-fold/*pltA^+^*: 8-fold mutant complemented with *pltA;* Δ8-fold/*ofaA^+^*: 8-fold mutant complemented with *ofaA;* Δ8-fold/*toxD^+^*: 8-fold mutant complemented with *toxD;* Δ8-fold/*pgnE^+^*: 8-fold mutant complemented with *pgnE;* Δ8-fold/*hcnB^+^*: 8-fold mutant complemented with *hcnB;* Picture is a representative of three replicates, and the experiment was repeated three times with similar results. The radius of the inhibitory zone represents the mean ± standard deviation of three replicates from a representative experiment, repeated twice. Different letters above the bars indicate that the data were statistically different (ANOVA, Tukey’s test, *p* < 0.05). **B.** Comparative genomics of wild type and Δ8-fold (LK892) of *P. protegens* Pf-5. Circular genome alignment highlights conserved regions and structural variations between the two strains. **C.** Genomic organization of the eight antibiotic loci in LK892. The mutations consist primarily of targeted deletions across multiple antibiotic biosynthetic clusters, with one locus disrupted by an insertion event, as indicated in the schematic. **D.** *D. pinodes* growth following exposure to Pf-5 and its derivative strains. Bacteria were inoculated on KB agar (200 ppm tetracycline) in a compartment spatially separated from PMA inoculated with the fungus. Plates were sealed and incubated at 28 °C for 14 days. Inhibitory activity was compared among WT, Δ*gacA*, Δ8-fold, and Δ8-fold/*hcnB^+^*strains. The change of picric acid paper to blue or purple indicates the production of hydrogen cyanide. Fungal colony diameter at 14 days post-inoculation is presented as bars representing the mean ± standard deviation of three replicates. **E.** Phenotypic characterization of the complemented strains. The eight antibiotic gene clusters were reintroduced into LK892 via plasmid-based complementation, and restoration of antibiotic-associated phenotypes was assessed using different approaches. Δ8-fold/*phlA*^+^ was tested via inhibition of *Aphanomyces euteiches.* Δ8-fold/*toxD*^+^ and Δ8-fold/*pgnE*^+^ were tested via inhibition of *Pseudomonas syringae.* Δ8-fold/*ofaA*^+^ was tested via a swarming motility assay on 0.6% agar M9 minimal medium. Δ8-fold/*hcnB*^+^ was tested via picric acid assay. Δ8-fold/*pltA*^+^ and Δ8-fold/*prnC*^+^ were cultured in nutrient broth supplemented with 1% glycerol and incubated for 24 h, and on pea meal agar for 7 days, respectively. Pyoluteorin and pyrrolnitrin were then extracted with ethyl acetate and visualized by TLC.

To identify the specific compounds involved, each deleted biosynthetic gene was individually complemented in the Δ8□fold mutant background, generating eight complemented strains that were evaluated for antagonistic activity against *D. pinodes*. Complementation of *rzxB* (rhizoxin biosynthesis) restored inhibition, generating the strain Δ8□fold/*rzxB* that clearly suppressed fungal growth (Fig. 5A). Similarly, complementation of *phlA* (DAPG biosynthesis) restored inhibition. Simultaneous complementation of both *rzxB* and *phlA* (Δ8□fold/*rzxB phlA*) also resulted in effective suppression of *D. pinodes*. Complementation of *prnC* (pyrrolnitrin biosynthesis) restored inhibitory activity as well, although the resulting inhibition zone was smaller than those observed for Δ8□fold/*phlA* and Δ8□fold/*rzxB*.

Among the Pf-5-derived antimicrobials, hydrogen cyanide (HCN) is a volatile compound. Restored HCN production of the complemented strain Δ8□fold/*hcnB* was confirmed via a colorimetric assay (Fig. 5D, 5E). The inhibition activity of Δ8□fold/*hcnB* was evaluated by co-culturing the bacterium with *D. pinodes* in a sealed plate assay. Results show that the colony size of *D. pinodes* was significantly decreased by co-culturing with the wild-type Pf-5 than the no-bacteria control (Fig. 5D), indicating Pf-5 produced volatile compound(s) to inhibit the fungus. The Δ*gacA* mutant and the Δ8□fold mutant had a less inhibition than the wild-type Pf-5. Importantly, strain Δ8□fold/*hcnB* restored the fungal inhibition to a level similar with the wild-type Pf-5.

Furthermore, individual complementation of the biosynthesis genes *toxD*, *ofaA*, *pgnE* and *pltA*, that are required for the production of toxoflavin, orfamide A, polyyne and pyoluteorin, respectively, failed to restore inhibition against *D. pinodes* (Fig. 5A). Successful restoration of these antimicrobial compounds in the complemented strains was confirmed via phenotypic assays (Fig. 5E).

Collectively, these results demonstrate that biosynthetic genes for rhizoxin, DAPG, pyrrolnitrin, and hydrogen cyanide are required for Pf□5-mediated inhibition of *D. pinodes*, indicating that these antimicrobial compounds contribute to pathogen suppression in cultures.

### 3.6 Rhizoxin and DAPG Contribute to the Biocontrol of *P. protegens* Pf-5 against *D. pinodes*

To investigate the roles of antimicrobial compounds in Pf□5-mediated biocontrol of pea AB, biocontrol efficacy was evaluated using Pf□5 and its derivative strains. Compared with the wild type Pf□5, both the Δ*gacA* and Δ8□fold mutants exhibited significantly reduced disease suppression, indicating that antimicrobial compounds regulated by GacA contribute to the biocontrol of pea AB.

Complementation of *rzxB* or *phlA* in the Δ8□fold mutant significantly enhanced biocontrol activity, as evidenced by reduced disease symptoms compared with plants treated with the Δ8□fold mutant alone (Fig. 6). Plants treated with the complemented strain Δ8□fold/*rzxB* showed no significant difference in DSI relative to those treated with wild type Pf□5, indicating full restoration of biocontrol activity. Complementation of *phlA* (Δ8□fold/*phlA*) increased disease suppression although the DSI was higher than in plants treated with wild type Pf□5, suggesting partial restoration of biocontrol efficacy. Similarly, the double complemented strain Δ8□fold/*rzxB phlA* exhibited biocontrol activity comparable to that of the wild type.

**Figure 6.**
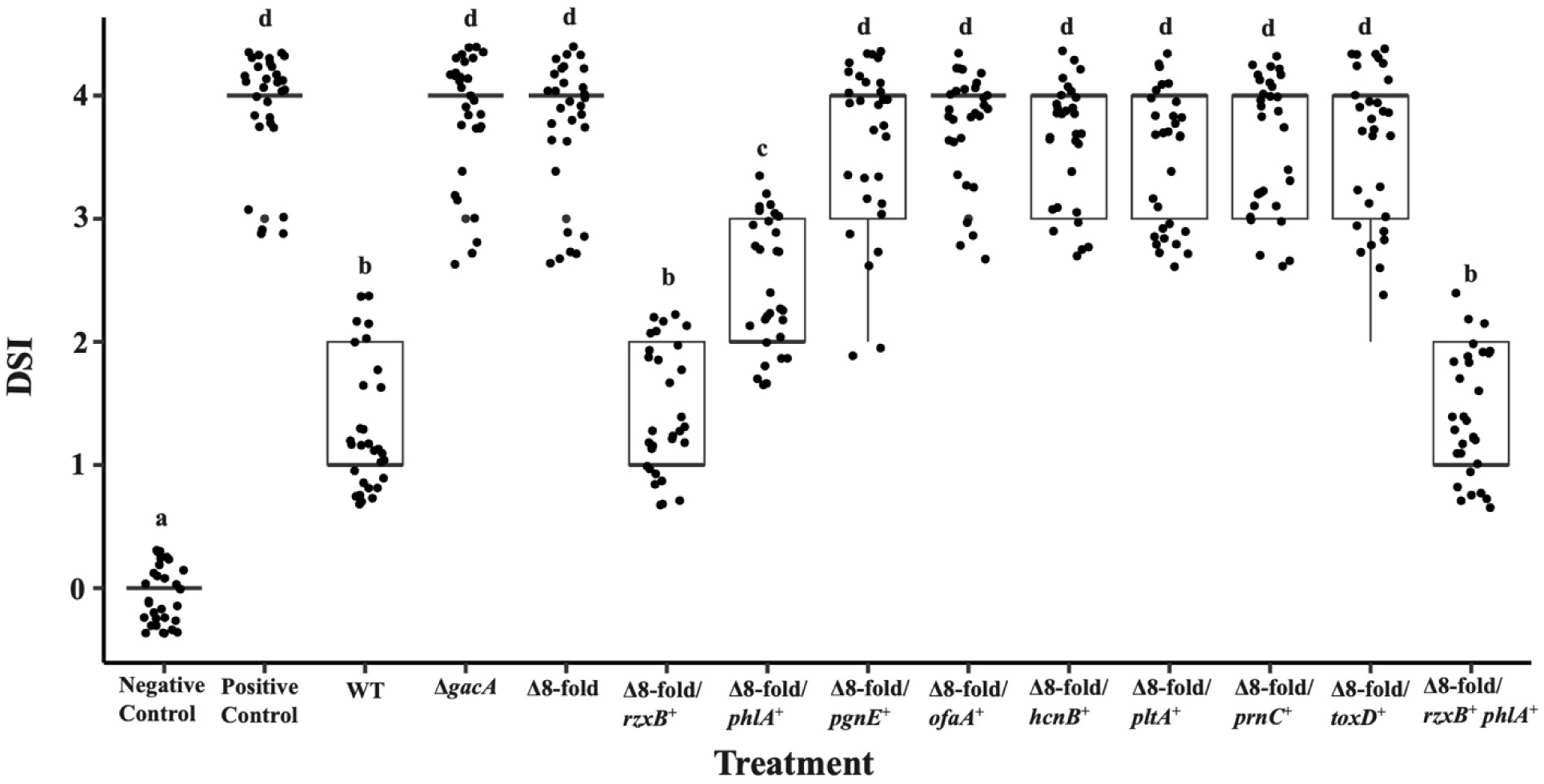
Biocontrol activity of *P. protegens* Pf-5 and its derivative strains against *D. pinodes* under greenhouse conditions. Disease severity index (DSI) of each treatment. Plants were treated with *P. protegens* Pf-5 and its derivative strains. WT: wild type; Δ8-fold: Δ*prnC*Δ*rzxB*Δ*pltA*Δ*hcnB*Δ*ofaA*Δ*phlA*Δ*pgnE*Δ*toxC*; Δ8-fold/*rzxB^+^*: 8-fold mutant complemented with *rzxB;* Δ8-fold/*rzxB^+^phlA^+^*: 8-fold mutant complemented with *rzxB* and *phlA;* Δ8-fold/*phlA^+^*: 8-fold mutant complemented with *phlA;* Δ8-fold/*prnC^+^*: 8-fold mutant complemented with *prnC;* Δ8-fold/*pltA^+^*: 8-fold mutant complemented with *pltA;* Δ8-fold/*hcnB^+^*: 8-fold mutant complemented with *hcnB;* Δ8-fold/*ofaA^+^*: 8-fold mutant complemented with *ofaA;* Δ8-fold/*toxD^+^*: 8-fold mutant complemented with *toxD;* Δ8-fold/*pgnE^+^*: 8-fold mutant complemented with *pgnE.* Data are presented as means ± standard deviation. Different letters indicate statistically significant differences among treatments (ANOVA, Tukey’s test, *P* < 0.05). The experiment was repeated independently three times, with 10 replicates per experimental set, yielding similar results.

Complementation of genes corresponding to the other six antimicrobial compounds in the Δ8□fold mutant did not restore biocontrol activity, indicating that these genes are not required for Pf□5-mediated control of pea AB.

Overall, these results demonstrate that *rzxB* and *phlA* are required for the biocontrol activity of Pf□5 against *D. pinodes*, suggesting that rhizoxin and DAPG contribute to the suppression of the pathogen *in planta*.

## 4. DISCUSSION

In this study, beneficial bacteria belonging to multiple genera, including *Pseudomonas*, *Bacillus*, *Paenibacillus*, and *Peribacillus*, were identified as effective suppressors of pea AB in greenhouse pot assays. Using *P. protegens* Pf□5 and derivative strains deficient in, or producing, specific antimicrobial compounds, we demonstrated that rhizoxin and DAPG are required for Pf□5 to inhibit *D. pinodes in vitro* and to suppress the AB pathogen on pea leaves *in planta*. To our knowledge, this is the first report demonstrating that rhizoxin produced by beneficial bacteria contributes to the biocontrol of a foliar disease.

Rhizoxin is a macrocyclic polyketide originally identified as a virulence factor of *Rhizopus microsporus* (formerly *Rhizopus chinensis*), the causal agent of rice seedling blight (Iwasaki et al. 1984). Subsequent work revealed that rhizoxin is produced by the endosymbiotic bacterium *Mycetohabitans rhizoxinica* (syn. *Burkholderia rhizoxinica*) rather than by the fungus itself (Partida Martinez & Hertweck 2005). Rhizoxin exhibits potent antimitotic activity against a broad range of eukaryotes, including plants (Iwasaki et al. 1984), fungi (Takahashi et al. 1990), and insects (Loper et al. 2016). Although rhizoxin has historically been studied as a virulence factor in pathogenic interactions, our results indicate that it can also function in beneficial plant-microbe interactions by suppressing fungal pathogens.

Consistent with our findings, *Pseudomonas* sp. Os17, a beneficial bacterium isolated from the rice rhizosphere, was reported to produce rhizoxin to control damping off caused by *Pythium ultimum* and root rot caused by *Fusarium oxysporum* (Takeuchi et al. 2015). Together, these studies suggest that microbial compounds traditionally regarded as virulence factors may also contribute to beneficial outcomes in plant-microbe interactions. Similarly, the bacterial type III secretion system, classically viewed as a virulence determinant mediating effector delivery by plant pathogens, has been shown to facilitate plant colonization by beneficial bacteria (Atanasković et al. 2025).

The observation that rhizoxin producing strains, including wild□ type Pf□5 and the complemented strain Δ8□fold/*rzxB□*, significantly reduced AB disease severity without causing visible phytotoxicity is particularly noteworthy. This finding suggests that Pf□5 produces rhizoxin on pea leaves at concentrations sufficient to inhibit the pathogen but below the threshold of plant toxicity. Alternatively, pea leaves may be relatively tolerant to rhizoxin produced by Pf□5. Supporting this possibility, a previous study reported that pea roots are less sensitive to Pf□5 derived rhizoxin than rice and cucumber roots (Loper et al. 2008).

In addition to rhizoxin, DAPG also appears to be produced by Pf□5 on pea leaves, as evidenced by the suppression of *D. pinodes* growth and reduced AB severity in plants treated with DAPG producing Pf□5 derivatives (e.g., Δ8□fold/*phlA*), but not with DAPG deficient strains (Fig. 6). Although DAPG producing bacteria are known to colonize leaf surfaces, the role of DAPG in the biocontrol of foliar diseases has been relatively understudied. Consistent with our results, DAPG was shown to be required for Pf□5 mediated suppression of leaf mold caused by *Botrytis cinerea* in cannabis (Balthazar et al. 2022).

DAPG is produced by many *Pseudomonas* species (Almario et al. 2017) and exhibits broad spectrum antimicrobial activity against a wide range of plant pathogens, including oomycetes (Keel et al. 1992; Pechy Tarr et al. 2007; Barahona et al. 2011; Lai et al. 2022), fungi (Raaijmakers & Weller 2001; Wei et al. 2004; Barahona et al. 2011; Balthazar et al. 2022), bacteria (Yan et al. 2017), and nematodes (Meyer et al. 2009). Research on DAPG has historically focused on rhizosphere processes and root disease management (Dobrzyński & Jakubowska 2025; Yang et al. 2025). Our study, together with previous work (Balthazar et al. 2022), supports a broader role for DAPG in the biocontrol of foliar diseases.

The observation that the DAPG□producing strain Δ8□fold/*phlA□* exhibited lower biocontrol efficacy than wild type Pf□5 and the rhizoxin producing strain Δ8□fold/rzxB (Fig. 6) suggests that rhizoxin is more effective than DAPG in controlling pea AB. This difference may reflect a higher sensitivity of *D. pinodes* to rhizoxin than to DAPG or differential production levels of these compounds on pea leaves. Rhizoxin and DAPG act through distinct mechanisms: rhizoxin binds *β□*tubulin and disrupts cell division (Takahashi et al. 1987), whereas DAPG compromises membrane integrity and induces reactive oxygen species production (Kwak et al. 2010). The combined use of two antimicrobial compounds with different modes of action likely enhances the robustness and consistency of disease suppression by Pf□5. Future studies should investigate whether rhizoxin and DAPG act additively or synergistically against *D. pinodes*.

Although the identified beneficial bacteria protected pea leaves from *D. pinodes* infection for up to 14 days, their biocontrol efficacy declined over time (Fig. 2). This decline was positively correlated with reduced bacterial populations on leaf surfaces, likely due to the harsh phyllosphere environment, which is nutrient poor, UV intensive, and characterized by rapid fluctuations in moisture and temperature (Meyer & Leveau 2011; Vorholt 2012). Similar declines in population size and biocontrol efficacy have been reported for *Bacillus subtilis* QST 713 in the control of tomato early blight (Abbasi & Weselowski 2014). Because foliar pathogens such as *D. pinodes* can infect host plants throughout the growing season, repeated application of biocontrol agents and integration with complementary disease management strategies that enhance bacterial persistence on leaves will likely be necessary for effective disease control (Wei et al. 2016).

Co□inoculation of beneficial bacteria did not enhance biocontrol efficacy against pea AB (Fig. 3). Similar outcomes have been reported in other foliar systems, where mixed bacterial inoculations often resulted in reduced disease suppression or diminished plant growth promotion compared with single strain applications (Saleem et al. 2017). Such reduced performance likely reflects incompatibility among strains due to competition for limited leaf nutrients and/or antagonistic interactions. These interactions can negatively affect bacterial growth and antibiotic biosynthesis, resulting in unchanged or even reduced disease control. In contrast, bacterial consortia frequently show additive or synergistic effects in root disease suppression (Liu et al. 2018; Solanki et al. 2019; Papp et al. 2021), possibly due to greater nutrient availability and spatial heterogeneity in the rhizosphere that promote functional complementarity among strains (Liu et al. 2023). Together, these findings underscore fundamental ecological differences between leaf and root environments that strongly influence microbial interactions and biocontrol outcomes.

Beyond antibiotic production, beneficial bacteria can suppress plant pathogens through additional mechanisms, including competition for nutrients and space (Spadaro & Droby 2016; Köhl et al. 2019), induction of plant defence responses (Pieterse et al. 2014), and promotion of plant growth (Etesami 2025; Lee et al. 2023). *Peribacillus* sp. LK832 did not inhibit *D. pinodes in vitro* (Fig. 4) but significantly reduced AB severity *in planta* (Fig. 1), suggesting that it suppresses disease through mechanisms other than direct antibiosis. Members of the genus *Peribacillus* are known to possess diverse traits relevant to agricultural applications (Manetsberger et al. 2023). For example, *Peribacillus butanolivorans* KJ40 suppressed cucumber anthracnose by inducing systemic resistance (Kim et al. 2023), while *P. frigoritolerans* strains have been shown to promote plant growth in *Arabidopsis thaliana* and canola (Marik et al. 2024; Swiatczak et al. 2024). Further research is needed to elucidate the mechanisms employed by *Peribacillus* sp. LK832 and other beneficial bacteria identified in this study to control pea Ascochyta blight.

## Acknowledgements

This study was supported by Specialty Crop Block Grant (SCBG) 25SC004910 from the Montana Department of Agriculture (MDA). We thank Kevin McPhee, Qian Wang, Amanda Stempke and Caden Kornak for their assistance in this work.

## Statement of No Conflict of Interest

The authors declare that they have no conflicts of interest related to the content of this manuscript.

## Data Availability

The data obtained in this study are available upon request or can be accessed with individual gene accession numbers of the identified beneficial bacteria as shown in the Figure 1E.

## Notes

### Competing Interest Statement

The authors have declared no competing interest.

